# Neutral genetic structuring of pathogen populations during rapid adaptation

**DOI:** 10.1101/2022.10.20.512995

**Authors:** Méline Saubin, Solenn Stoeckel, Aurélien Tellier, Fabien Halkett

**Affiliations:** Université de Lorraine, INRAE, IAM, F-54000 Nancy, France; Professorship for Population Genetics, Technical University of Munich, Freising, Germany; INRAE, Agrocampus Ouest, Université de Rennes, IGEPP, F-35653 Le Rheu, France; DECOD (Ecosystem Dynamics and Sustainability), INRAE, Institut Agro, IFREMER, 35042, Rennes, France

**Author notes:** Corresponding author: Fabien Halkett INRAE Centre Grand-Est – Nancy, UMR 1136 Interactions Arbres-Microorganismes, F-54280, Champenoux, France.

**Keywords:** Forward demogenetic model, Plant pathogen, Host adaptation, Complex life cycles, Time-series clustering, Population genetic structure

## Abstract

Pathogen species are experiencing strong joint demographic and selective events, especially when they adapt to a new host, for example through overcoming plant resistance. Stochasticity in the founding event and the associated demographic variations hinder our understanding of the expected evolutionary trajectories and the genetic structure emerging at both neutral and selected loci. What would be the typical genetic signatures of such a rapid adaptation event is not elucidated. Here, we build a demogenetic model to monitor pathogen population dynamics and genetic evolution on two host compartments (susceptible and resistant). We design our model to fit two plant pathogen life cycles, ‘with’ and ‘without’ host alternation. Our aim is to draw a typology of eco-evolutionary dynamics. Using time-series clustering, we identify three main scenarios: 1) small variations in the pathogen population size and small changes in genetic structure, 2) a strong founder event on the resistant host that in turn leads to the emergence of genetic structure on the susceptible host, and 3) evolutionary rescue that results in a strong founder event on the resistant host, preceded by a bottleneck on the susceptible host. We pinpoint differences between life cycles with notably more evolutionary rescue ‘with’ host alternation. Beyond the selective event itself, the demographic trajectory imposes specific changes in the genetic structure of the pathogen population. Most of these genetic changes are transient, with a signature of resistance overcoming that vanishes within a few years only. Considering time-series is therefore of utmost importance to accurately decipher pathogen evolution.

## 1 Introduction

Pathogen populations commonly endure large demographic variations, including repeated bottlenecks and founder events (McDonald, 2004; Barrett et al., 2008). These are often associated with selective events, with the adaptation of pathogens to their hosts, that sets resource availability over time and space (Stukenbrock and McDonald, 2008). Yet, we have limited theoretical knowledge of how such events shape the evolutionary trajectories of pathogens and what would be the typical genetic signatures of the interplay of strong demographic and selective events on the pathogen population.

By contrast, the population genetic structures of pathogen species has been extensively investigated empirically (see for reviews McDonald and Linde, 2002; Gladieux et al., 2011; Möller and Stukenbrock, 2017; Hessenauer et al., 2021). The apportionment of genetic variability is most often examined through space and between hosts with the aim to provide insights on the route of migration, the extent of dispersal and the delineation of host-specific populations. Focusing on host adaptation, these investigations highlighted a wide array of patterns that range from strong genetic structuring that last for decades despite large gene flow (Leroy et al., 2013; Susi et al., 2020) to the lack of genetic differentiation despite evidence for host adaptation (Linde et al., 2002; Travadon et al., 2011; Siah et al., 2018).

Some pathogen species can also reveal a transient population genetic structure, with marked population differentiation that vanishes over few years (Persoons et al., 2017). More specifically, the same pathogen species can display contrasted genetic structures in different environments (Halkett et al., 2010). This points to the importance of demographic events in the emergence of genetic structures. Yet theoretical population genetics classically assumes demographic equilibrium or simplistic demographic scenario to build predictions. Moreover, understanding the emergence of a genetic structure requires deciphering the temporal evolution of population genetic indices, which is rarely done both theoretically and empirically (Saubin et al., 2023b). Finally, the stochastic nature of the evolution of genetic diversity and structuring of populations blurs and even hinders our comprehension of their expected dynamics. We thus need *ad hoc* approaches to identify and quantify the different types of evolutionary trajectories and how they translate into different genetic structures, especially for species under management plan.

Most pathogens have complex life cycles (Agrios, 2005), often exhibiting mixed reproductive systems and partial clonality. We can distinguish autoecious pathogens, which complete their life cycle on a unique host species, from heteroecious pathogens which need two different and successive host species to complete their life cycle (Moran, 1992; Lorrain et al., 2019). Population genetics can be used to describe the neutral genetic signatures and evolution of sexual populations, but the partial clonality of such species makes the study of these genetic signatures much more complex (Orive, 1993).

The lack of theoretical developments dedicated to understand the emergence of genetic structure in pathogens prompts us to develop a new demogenetic model (see a definition of such models in Lamarins et al., 2022). Coupling epidemiology and population genetics provides insights into the mechanisms underpinning pathogen evolution acting at both short (ecological) and long (evolutionary) time scales (Milgroom and Peever, 2003; Archie et al., 2009). As such, it enables the study of genetic signatures of strong and rapid selective events (Saubin et al., 2023a). The interplay between demography and selection is captured by monitoring both selected and neutral loci. It allows in particular detailed analyses of transition periods (Day and Proulx, 2004; Day and Gandon, 2007; Bolker et al., 2010), through variables like the pathogen population size, affecting both the disease incidence in epidemiology and the impact of genetic drift in population genetics (McDonald, 2004; Z^̌^ivkovíc et al., 2019).

In this article, we focus on pathogen adaptation to its host as a case study to delineate the different scenarios of evolutionary trajectory that can occur during the same adaptive event. Two main qualitative mechanisms by which pathogens adapt to their hosts are usually considered: the matching allele and the gene-for-gene model (Agrawal and Lively, 2002; Thrall et al., 2016). In this study, we focus on the genefor-gene model, as it accounts for most plant-pathogen interactions (Thrall et al., 2016) and attracts a great deal of breeding efforts because, in most cases, it confers complete host immunity. According to the genefor-gene model, genetic resistance prevents infection from a class of pathogen genotypes called *avirulent*. In agrosystems, the deployment of pure resistant plants exerts a strong selection pressure on the pathogen population, that favours any variant that can infect the resistant host (Zhan et al., 2015). This class of pathogen genotypes is called *virulent*. The infection success is determined by a single locus, with avirulent and virulent alleles. The spread of virulent individuals on resistant hosts leads to a so-called resistance overcoming event, which can result in severe epidemics (Johnson (1984); Pink and Puddephat (1999); Brown and Tellier (2011); Burdon et al. (2016)), and in rapid and drastic demographic changes for the pathogen population (Persoons et al., 2017; Saubin et al., 2021). In our model, hosts are considered as static compartments because we assume that infections do not lead to hosts’ death, and the generation time of the pathogen is much shorter than that of the hosts. We assume the simplest case of two host compartments: susceptible hosts can be infected by all pathogen genotypes while resistant hosts can only be infected by virulent individuals (*i.e.* individuals with only the virulent allele at the avirulence locus).

We model pathogen population dynamics and genetic evolution to investigate the impact of the pathogen life cycle on these selective and demographic dynamics using a demogenetic approach, tracking the exact evolutionary trajectories forward in time. We perform simulations under several realistic scenarios of resistance overcoming. We build a random simulation design to ensure all types of events are covered. Then, we use a clustering method dedicated to time-series variations applied to the temporal change of neutral population genetic indices to identify the main scenarios of eco-evolutionary dynamics. We conclude by commenting on the typology of these dynamics and the potential to use our simulation framework to analyse real datasets.

## 2 Materials and methods

### 2.1 Model description

We develop an individual-based, compartmental and forward in time demogenetic model. It couples population dynamics and population genetics to follow through time the exact evolutionary trajectory of different genotypes at a selected locus and at neutral genetic markers scattered in the genome. The model is similar to the model described in Saubin et al. (2021) and Saubin et al. (2023b), but its treatment differs. Here we focus on the expectations, in terms of neutral population genetics, when varying the five main input parameters (Table 1). A model overview is provided in Figure 1. Descriptions of the reproduction and migration events are provided in Appendix A.1 and Appendix A.2.

**Figure 1:**
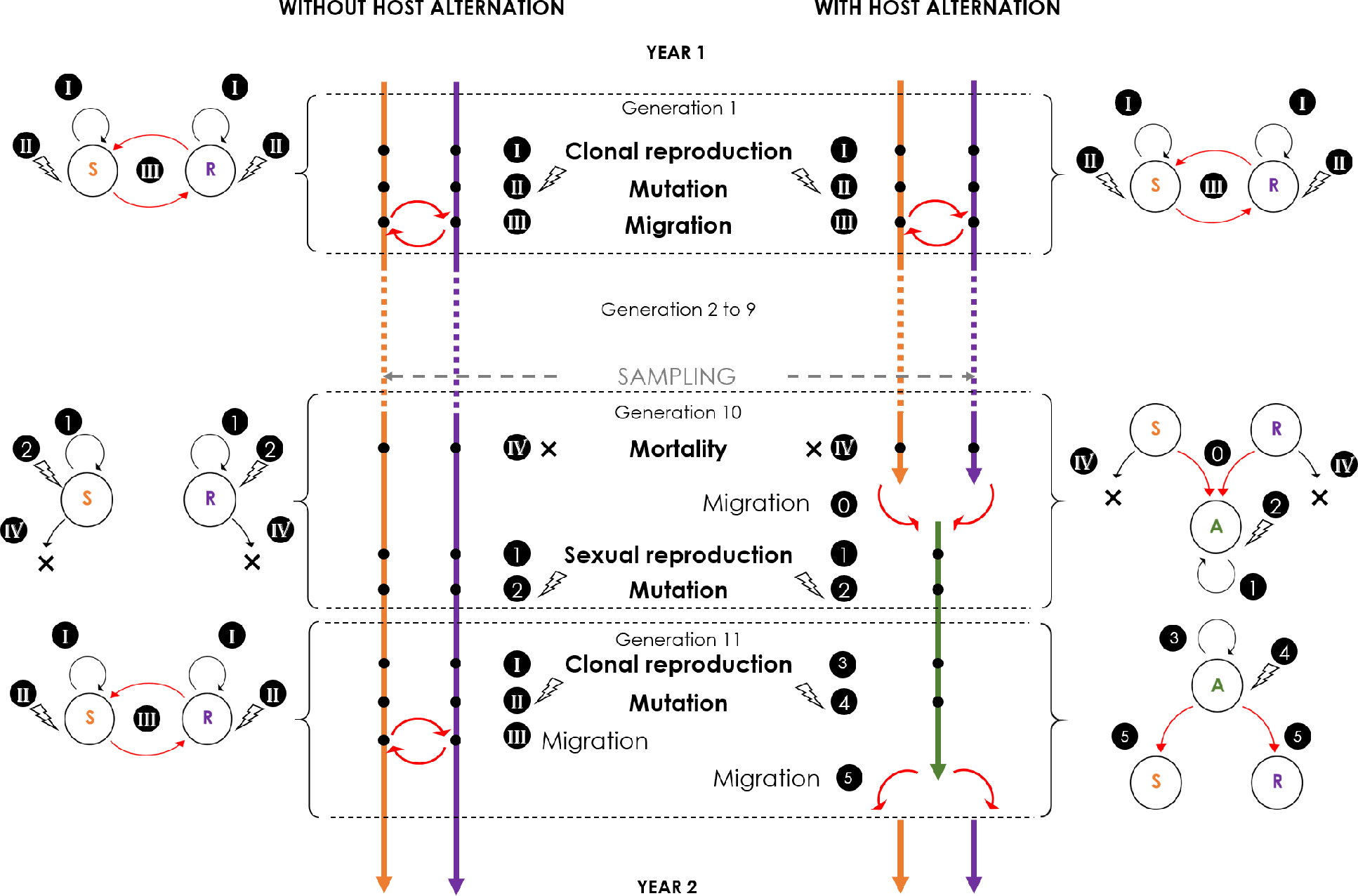
Modelling steps for each simulated year with the three S, R, and A compartments (adapted from Saubin et al., 2023b, Figure 1). Each year is composed of *g* = 11 generations. During the clonal phase (generation 1 to *g −* 2), each generation is composed of three steps identical between both life cycles: (I) clonal reproduction; (II) migration of a proportion *m* of each population between R and S; (III) mutation at all neutral markers with a mutation rate *µ*. At the end of the clonal phase, the pathogen overwinter as a dormant stage and is subjected to (IV) mortality of a proportion *τ* of each population. Then, the sexual phase (generation 10) differs depending on the life cycle: (0) represents the migration of all individuals from R and S towards A; (1) sexual reproduction; (2) mutation of all neutral markers with a mutation rate *µ*. This sexual phase is followed by a new clonal phase, which is identical ‘without’ alternation to the first clonal phase and ‘with’ alternation: (3) represents the clonal reproduction; (4) mutation of all neutral markers with a mutation rate *µ*; (5) migration of all individuals from A towards R and S. A sampling takes place every year at the end of generation 9 on S and R.

**Table 1:**
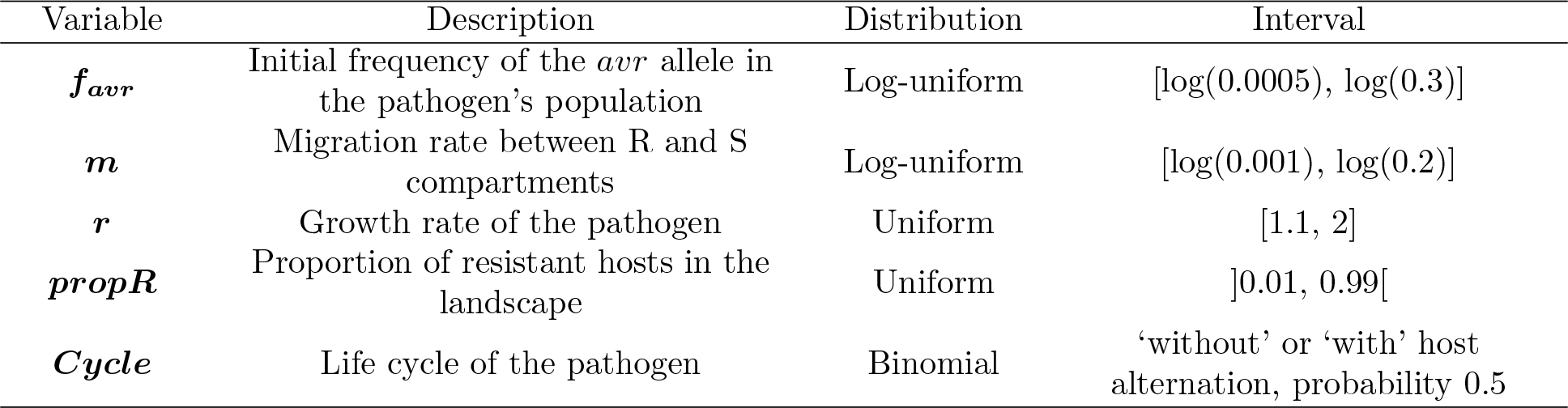
Input parameters and their range of variations for the random simulation design. Each simulation is run for 400 generations.

The model simulates the evolution over time of a population of diploid pathogens. Pathogen life cycles usually include several generations (*i.e.* infections of the same host species or not) that consist in successive steps of within host growth, clonal or sexual reproduction and spread. We consider life cycles commonly found in temperate pathogen species, with seasonal variation in reproductive mode. These pathogens switch from several generations of clonal reproduction during the epidemic phase to sexual reproduction once a year, in winter (Agrios, 2005). This model is designed to simulate two distinct pathogen life cycles: ‘with’ or ‘without’ host alternation for the sexual reproduction (Boolean parameter *Cycle*). During the clonal phase, the life cycles are similar and the pathogen evolve on two host compartments: susceptible (S) and resistant (R). During the sexual phase, the life cycles differ: ‘with’ alternation, pathogens have to migrate to an alternate host (A) to perform sexual reproduction. ‘Without’ alternation, pathogens stay on R and S compartments for the sexual reproduction (A remains empty). Thereafter, when we refer to the pathogen life cycle, we refer specifically to the presence or absence of an host alternation during the sexual reproduction, the rest of the life cycle being otherwise identical.

We do not consider spatial substructure among compartments. We assume fixed carrying capacities of pathogens for each host compartment, *K_R_*, *K_S_* and *K_A_* for compartments R, S and A respectively. They represent the maximum amount of pathogens that each host compartment can sustain. We thus consider each host compartment to be ‘static’. This assumption holds as long as the pathogen generation time is considered much shorter than that of the hosts, and the pathogen does not kill its host. It is the case for example for biotrophic pathogens of perennial plants, such as grape-wine mildew or poplar rust pathogens.

We consider that a year consists of *g* = 11 generations: *g−*1 rounds of clonal multiplication plus one sexual reproduction event. This corresponds to the expected generation time of the fungal pathogen responsible for the poplar rust disease (Hacquard et al., 2011). Three basic steps are modelled at each clonal generation: reproduction following a logistic growth (with growth rate (*r*) and carrying capacity *K_R_* or *K_S_* depending on the compartment considered, see Appendix A.1), mutation of neutral loci (at a fixed mutation rate *µ*, see below), and a two-way migration (migration rate *m*, see Appendix A.2), from S to R and vice versa (Appendix A, Figure 1). At the end of clonal multiplication, random mortality is applied to the pathogen population (at rate *τ*) because some individuals fail to overwinter. Then, sexual reproduction occurs. It differs between life cycles, considering or not the obligate migration to the alternate host before mating. For the life cycle ‘with’ alternation, the generation of sexual reproduction is followed by one generation of clonal multiplication on A before the pathogen emigration to S and R.

Following the gene-for-gene model, we consider the very simple genetic architecture for pathogen adaptation to the resistant host with a single bi-allelic avirulence locus: a dominant avirulent allele (*Avr*) and a recessive virulent allele (*avr*). All individuals (genotypes *Avr*/*Avr*, *Avr*/*avr* and *avr*/*avr*) survive on S and A, while only individuals with the homozygous genotype *avr*/*avr* (called virulent individuals) can survive on R. We assume no fitness cost of virulence because fitness costs are not systematic in plant pathogens and key to drive coevolution scenarios (see Leach et al., 2001; Brown and Tellier, 2011 for reviews). We consider that evolution stems from standing genetic variation, with an initial frequency of the *avr* allele (*f_avr_*) set after the burn-in phase (see Section 2.2.2 below), and we do not consider mutation at the avirulence locus. In addition to the avirulence locus, we simulate the evolution of 100 independent neutral genetic markers with a mutation rate *µ* = 10*^−^*^3^. Each locus has four possible allelic states under a classical k-allele mutation model (Wright, 1949). Upon mutation, an allele changes into any of the three other allelic states with equal probability. At these 100 loci, we compute yearly 10 classical population genetic indices before the sexual reproduction on both R and S compartments (Table 2, and see Section 2.2.2 for a more complete description). These indices are chosen to 1) describe intra-population genetic and genotypic diversity, 2) measure overall linkage disequilibrium, and 3) assess genetic differentiation (*F_ST_*). The differentiation is considered between populations on R and S at a given generation, and through time between a population on R or S at a given generation and the initial genotypic state. The variation in these indices captures the footprints of the different processes expected to occur during resistance overcoming: effects of reproductive mode, founder event followed by expansion, coancestry and admixture between R and S.

**Table 2:**
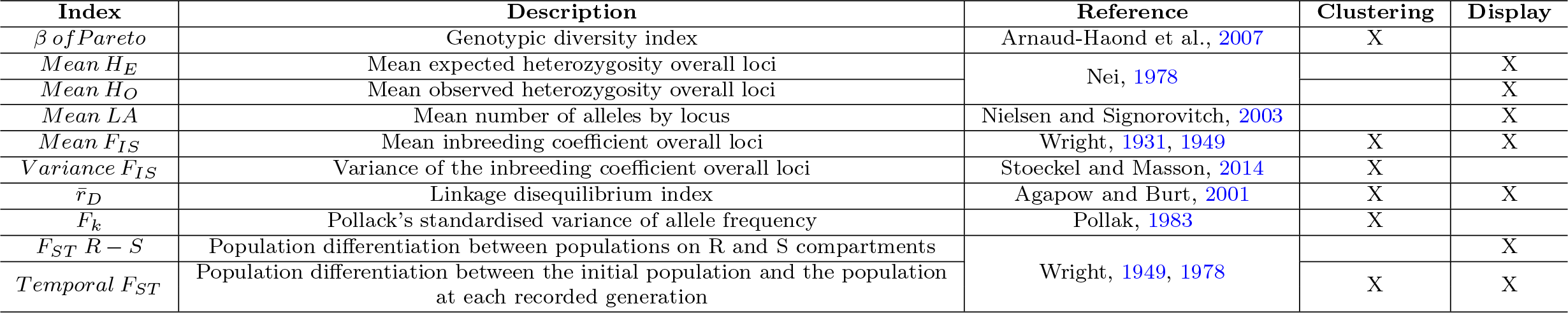
Description of population genetic indices computed in the model and calculated each year before the sexual reproduction on both R and S compartments. The ‘Clustering’ column represents the indices used for clustering analyses, the ‘Display’ column represents the indices whose evolution is presented in graphical form.

| Index | Description | Reference | Clustering | Display |
| --- | --- | --- | --- | --- |
| $\beta$ of *Pareto* | Genotypic diversity index | Arnaud-Haond et al., 2007 | X | |
| $Mean H_E$ | Mean expected heterozygosity overall loci | Nei, 1978 | | X |
| $Mean H_O$ | Mean observed heterozygosity overall loci | | | X |
| $Mean L_A$ | Mean number of alleles by locus | Nielsen and Signorovitch, 2003 | | X |
| $Mean F_{IS}$ | Mean inbreeding coefficient overall loci | Wright, 1931, 1949 | X | X |
| $Variance F_{IS}$ | Variance of the inbreeding coefficient overall loci | Stoeckel and Masson, 2014 | X | |
| $\bar{r}_D$ | Linkage disequilibrium index | Agapow and Burt, 2001 | X | X |
| $F_k$ | Pollack's standardised variance of allele frequency | Pollak, 1983 | X | |
| $F_{ST} R - S$ | Population differentiation between populations on R and S compartments | Wright, 1949, 1978 | | X |
| $Temporal F_{ST}$ | Population differentiation between the initial population and the population at each recorded generation | | X | X |

**Table 3:**
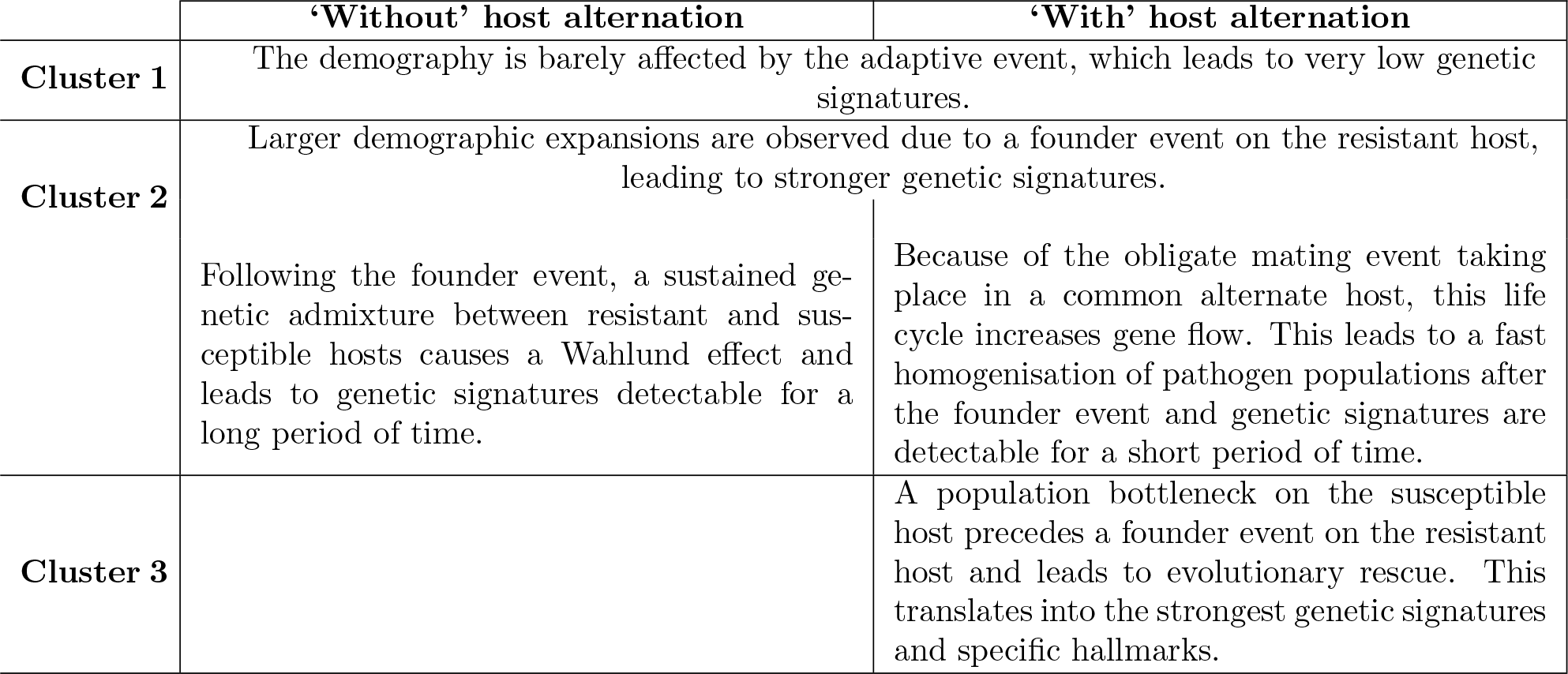
Summary of the typology of eco-evolutionary dynamics for each cluster, depending on the considered life cycle.

### 2.2 Simulations and temporal dynamic analyses

#### 2.2.1 Method overview

The model allows us to investigate the genetic consequences of rapid adaptation. A general overview of the method and the successive steps are presented in Figure S1. First, we build a random simulation design by drawing randomly five input parameters in defined distributions (Table 1). For each independent parameter combination, we simulate forward in time the stochastic evolutionary trajectory and track the population state at each sampled generation by computing ten classical population genetic indices (output trajectories). We then retain only simulations leading to population adaptation. We regroup simulations leading to similar population genetic evolution using a classical time-series clustering approach. By deriving mean dynamics of the clustered time-series (centroids), we aimed at drawing a typology of the main eco-evolutionary dynamics, without exhaustively analysing each individual trajectory. To avoid redundancy of information, we base the clustering on the output trajectories from six population genetic indices that are orthogonal by their mathematical construction. Finally, we compare clusters through graphical and sensitivity analyses and identify the main scenarios of eco-evolutionary dynamics.

#### 2.2.2 Model implementation and simulations

The model is implemented in Python (version 3.7, van Rossum, 1995) and Numpy (Harris et al., 2020). Each simulation starts with genotypes randomly drawn from the four possible alleles followed by a burn-in period of 11,000 generations under a constant population size of *K_S_* individuals. In this way, we ensure that the pathogen population is at the mutation-drift equilibrium before overcoming the resistance. At the avirulence locus under selection, a proportion *f_avr_* of virulent alleles is introduced randomly (replacing avirulent *Avr* alleles) on S after the burn-in period as initial standing genetic variation. Homozygous *avr*/*avr* and heterozygous *Avr*/*avr* individuals can therefore be initially present, based on the frequency *f_avr_*. Simulations are run with a fixed total carrying capacity for the host population sizes of each host species, *K* = *K_A_*= *K_R_* + *K_S_*= 10,000. We define *propR* as the proportion of resistant hosts in the cultivated landscape 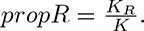

We run a random design set of 30,000 independent simulations. Each combination of parameter values is drawn at random in defined prior distributions (Table 1). Each simulation is run for 400 generations (with 11 generations per year), which amounts to 36 years. During this period, nearly all replicates reach new steady states including the settlement on R and loss of the avirulent *Avr* allele or the extinction of the pathogen population. To focus on the genetic signatures of a resistance overcoming, only the simulations with at least 60% of virulent *avr* alleles at the end of the simulation are kept for this analysis. This threshold is chosen to focus on resistance overcoming events, by ensuring that the settlement on the resistant compartment does occur during the simulated period.

To track the genetic dynamics of populations, we computed the temporal variation of ten population genetic indices listed Table 2, classically used to assess evolutionary forces, including temporal changes in demography, reproductive modes and adaptation (*e.g.* Allen and Lynch, 2012; Skoglund et al., 2014; Arnaud-Haond et al., 2020). The F-statistics were tracked to quantify the level of apportionment of genetic variability within and between populations sampled over time or over different compartments (Wright, 1931, 1949, 1978). *β of Pareto* accounting for genotypic diversity (Arnaud-Haond et al., 2007), *r̄_D_* accounting for overall linkage disequilibrium (Agapow and Burt, 2001), mean and variance over loci of *F_IS_* (inbreeding coefficient) accounting for the proportion of the genetic variance contained within individuals were tracked to understand the importance of clonal reproduction to contribute to the population dynamics (Halkett et al., 2005; Stoeckel et al., 2021). We also tracked observed *H_O_* and expected *H_E_* gene diversity as well as the mean number of alleles *Mean LA* for their different sensitivities to a bottleneck (Luikart et al., 1998). Finally, we calculated the population size estimator *F_k_* based on time-step changes in allele frequencies (Pollak, 1983).

#### 2.2.3 Comparisons of temporal dynamics

Analyses of changes in population genetic indices are performed using the R statistic software (Team, 2018). We present all results on the S compartment because it enables to compare the effect of life cycles, all else being equal, and to represent the genetic signatures expected without selection. When needed, we refer to the evolution of indices on the R compartment provided in supplementary data.

To analyse the dynamics of population genetic indices from the random simulation design, we performed hierarchical agglomerative clustering analyses regrouping simulations with similar dynamics (*i.e.* temporal evolutionary trajectories on S) using classical *Dynamic Time Warping* distance. Distinct clustering analyses are performed for the two life cycles using the package Dtwclust (Sardá-Espinosa, 2019), dedicated to the clustering of time-series. To avoid redundancy in genetic information, clustering analyses are based on the temporal dynamics of the six indices that are orthogonal by their mathematical construction (*i.e.* with the least mathematical redundancy among them): *β of Pareto* for genotypic diversity, *r̄_D_* for overall linkage disequilibrium between loci, *Mean F_IS_* and *V ariance F_IS_* for allele identity within individuals, *F_k_* for the variation of allele identity between individuals within a compartment, and *Temporal F_ST_* for the genetic differentiation (between time points on S). We perform a multivariate analysis by concatenating in time the temporal dynamics of the six normalised indices. We then build a distance matrix between all simulations based on the distance ‘DTW basic’, a classical *Dynamic Time Warping* distance dedicated to the measure of similarity between two temporal sequences. We use the norm ‘Euclidean distance’ for the local cost matrix to accentuate the distance between the most discrepant simulations and the step pattern ‘symmetric2’ (which is one of the common transition types and is normalizable, symmetric, with no local slope constraints). We perform hierarchical clustering based on the distance matrix, with the agglomeration method ‘Ward.D2’, which minimises within-cluster variance and combines clusters according to their smallest squared dissimilarities. We compare hierarchical clustering for a number of clusters ranging from two to eight. For each life cycle, we select the number of clusters that maximises the Calinski-Harabasz index, calculated as the ratio of the inter-cluster variance and the sum of intra-cluster variances (Arbelaitz et al., 2013).

To understand how input parameters impact simulation clustering, we represent the distribution of input parameters for each cluster and both life cycles. We assess significant differences between distributions of parameter values using pairwise Kruskal-Wallis tests. To rank the impact of input parameters on the clustering, we perform a dominance analysis for each life cycle and represent the estimated general dominance of each parameter on cluster assignments. Dominance analyses are performed with the package Domir (Luchman, 2022).

For each cluster, we illustrate the evolution of population genetic indices through time. For the sake of clarity and to limit the amount of information, we focus in this article on the most informative four intrapopulation and two inter-population indices: *r̄_D_*, *Mean F_IS_*, *Mean H_E_*, *Mean LA*, *Temporal F_ST_*, *F_ST_ R − S* (Table 2). We display for each cluster and both life cycles the mean and standard error of these six population genetic indices calculated at each generation over all simulations assigned to a given cluster.

To illustrate the realised dynamics of each population genetic index, we complement these results by displaying a representative realised simulation of each cluster. We choose for that the medoid, that is the simulation that minimises the average distance to all other simulations in the same cluster. To highlight the effect of selection, we supplement each representation of the medoid dynamics with the corresponding null model dynamics, that is a simulation run under the same set of parameter values but without selection (*f_avr_* = 0). The deviation between medoid and null model dynamics highlights the specific signatures of selection. To interpret genetic changes with respect to resistance overcoming, we calculate the generation at which pathogens overcome resistance as the generation for which 1% of R is occupied by virulent individuals for the first time in a simulation. In addition, we define the generation of settlement as the first generation in which a virulent individual migrates to R.

Some simulations lead to evolutionary rescue, that is the settlement on R resulting in the recovery of the population collapse on S and preventing population extinction. To understand which input parameter combinations favour the occurrence of evolutionary rescue, we calculate for each simulation the growth rate threshold under which the population goes extinct if R is not accessible (population extinction in the ‘null model’). We thus obtain the proportions of evolutionary rescue events in each cluster and perform a Fisher’s exact test to assess the significance of the assignments to different clusters.

## 3 Results

### 3.1 Influence of model parameters on the evolutionary dynamics

The clustering based on the random simulation design results in a partition into two clusters ‘without’ host alternation and into three clusters ‘with’ host alternation. For both life cycles, we name Cluster 1 the cluster that regroups the majority of the simulations (72 % and 78 % of the simulations ‘without’ and ‘with’ host alternation, respectively, Table S3). Cluster 1 displays small genetic changes through time for nearly all indices (Figure 2). Conversely, the other clusters display stronger changes in population genetic indices (Figures 2, 3).

**Figure 2:**
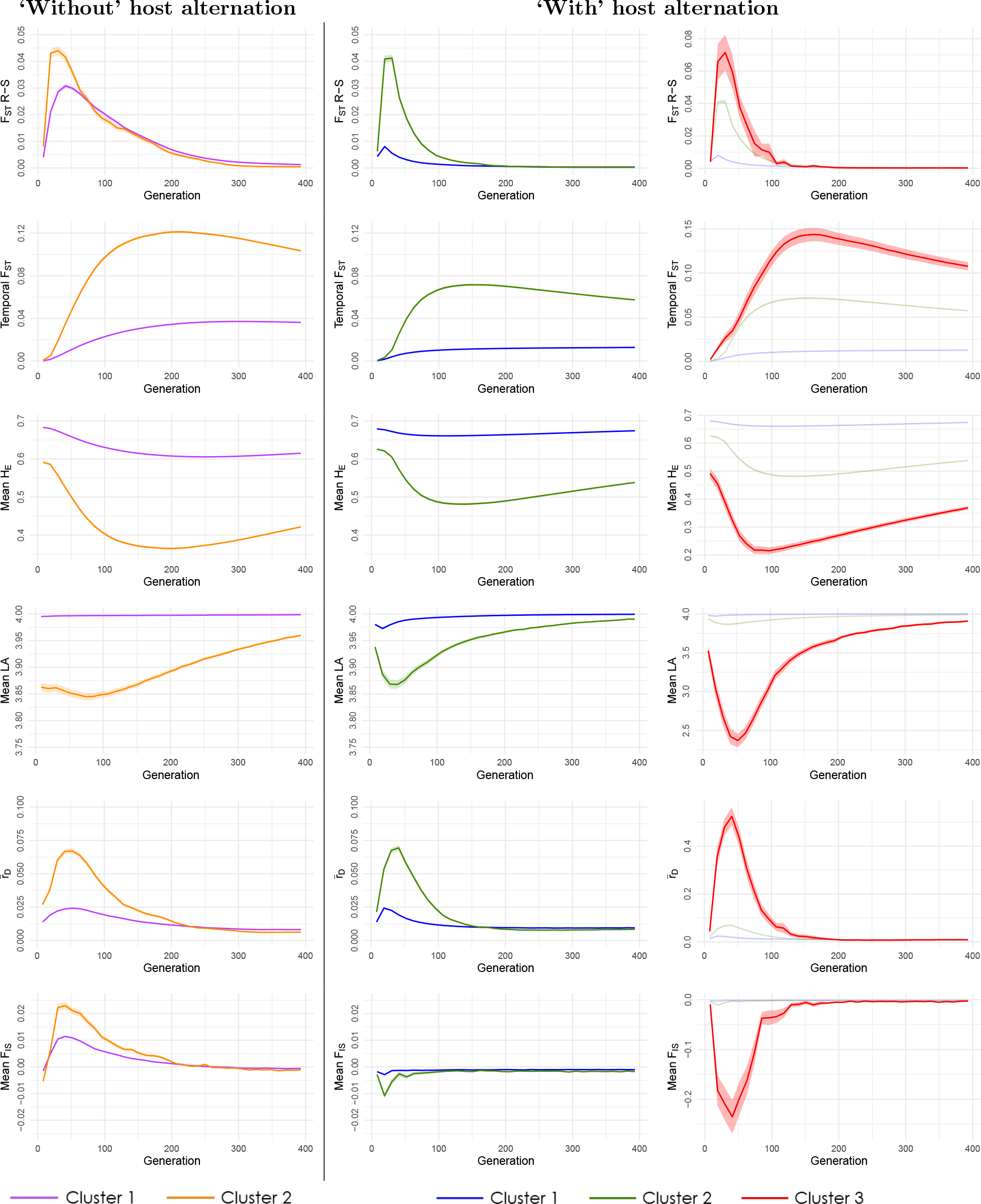
Temporal evolution of population genetic indices. The plotted results correspond to the mean temporal dynamics for all simulations in the corresponding cluster, among all simulations of the random simulation design. Populations are sampled on S, except for *F_ST_ R − S*. Shaded colour bands correspond to standard error intervals. For clarity, we apply the same scale for Cluster 1 and Cluster 2 of both life cycles, but we use a different scale to display the changes in population genetic indices for Cluster 3 ‘with’ host alternation.

**Figure 3:**
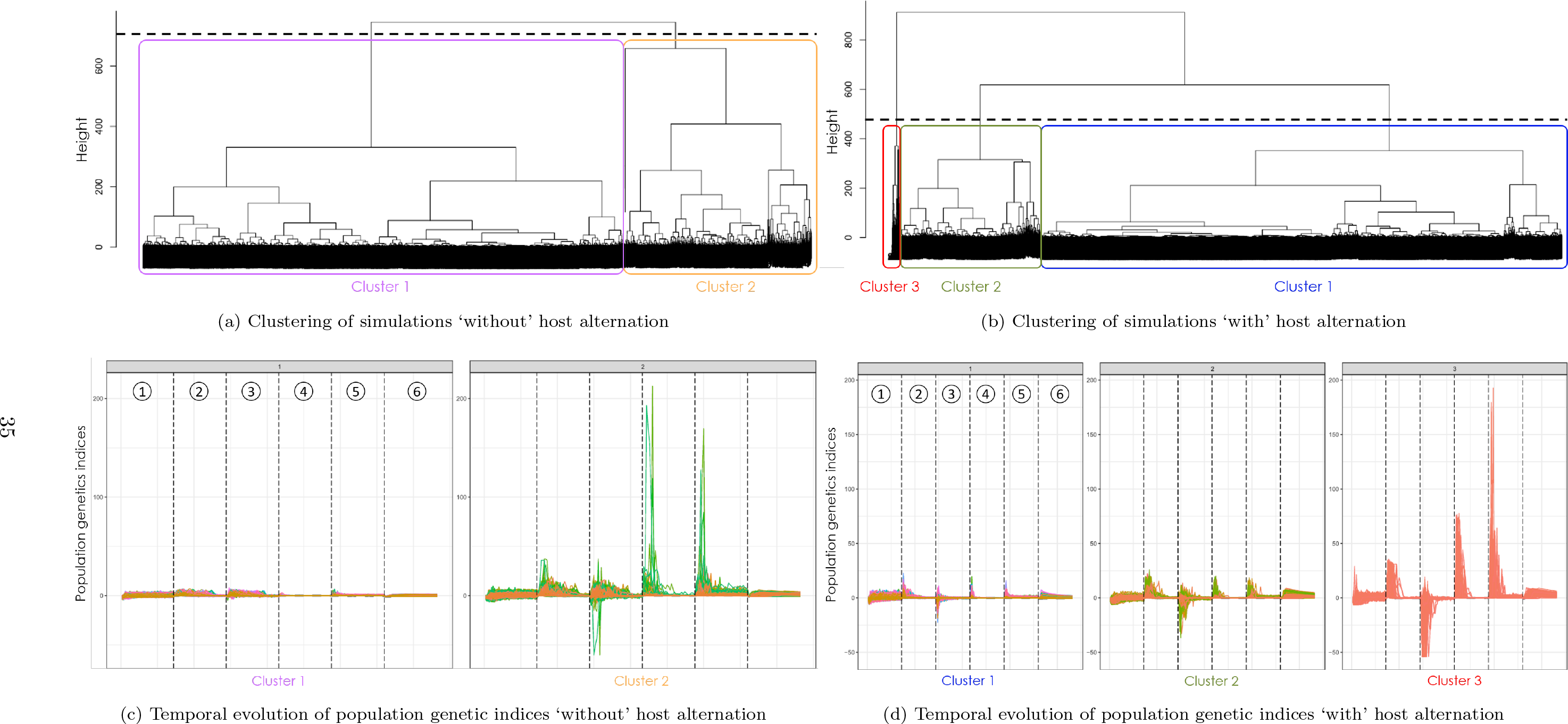
Clustering of simulations ‘without’ (a, c) and ‘with’ (b, d) host alternation. The clustering analyses are based on the temporal evolution of six population genetic indices. (a, b) represent the dendrograms obtained from the clustering analyses. Distances between simulations were calculated based on the dynamic time warping distance, with the norm ‘Euclidean distance’ and the step pattern ‘symmetric 2’. (c, d) represent for each cluster the concatenation of the temporal evolution of six normalised populations genetics indices, in the following order: (1) *β of Pareto*, (2) *r̄_D_*, (3) *Mean F_IS_*, (4) *V ariance F_IS_*, (5) *F_k_*, (6) *Temporal F_ST_* .

To understand the origin of the different types of dynamics, we jointly examine the influence of the input parameters on cluster delineation (Figure S2) and the difference in the distributions of parameter values for each cluster (Figure 4).

**Figure 4:**
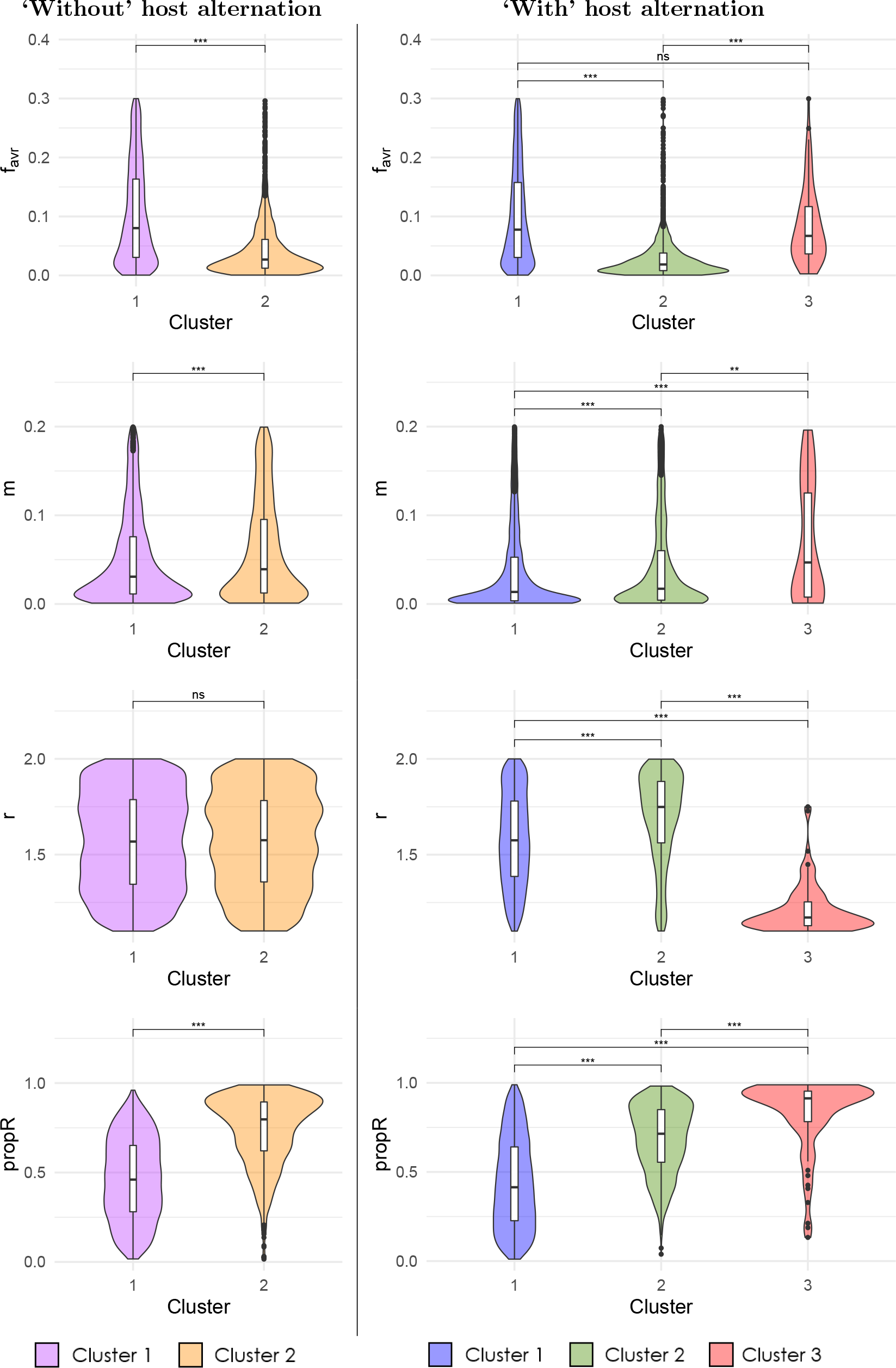
Distribution of epidemic parameters for each cluster obtained from the dynamics of population genetic indices. Differences between distributions were statistically assessed using pairwise Kruskal-Wallis tests: ns, non-significant; ·, *P −value <* 0.1; *, *P −value <* 0.05; **, *P −value <* 0.01; ***, *P −value <* 0.001. ‘Without’ host alternation, Cluster 1 and Cluster 2 are composed of 4067 and 1575 simulations, respectively. ‘With’ host alternation, Cluster 1, Cluster 2 and Cluster 3 are composed of 5858, 1581, and 81 simulations, respectively.

‘Without’ host alternation, the most influential parameter for cluster delineation is the proportion of resistant hosts (*propR*), followed by the initial frequency of avirulent allele (*f_avr_*), and, to a lesser extent, the migration rate (*m*) (Figure S2). As such, Cluster 2 is composed of simulations with significantly larger values of *propR* and significantly smaller values of *f_avr_* and slightly, albeit significantly, lower *m* (Figure 4). The growth rate (*r*) has no effect on cluster delineation (and consistently there is no significant difference in parameter distribution across clusters). It is important, however, to keep in mind that the analysis only considers simulations where R is overcome by the pathogen.

‘With’ host alternation, we observe the same differences between Cluster 1 and Cluster 2, with similar variations in the distribution of parameter values and ranking of parameter effects. The only difference is that *r* has an effect on cluster delineation with significantly higher growth rates for Cluster 2. Cluster 3 represents a particular case with skewed distributions towards very high values of *propR* and low values of *r* (Figure 4). Cluster 3 also displays high values of *f_avr_* and higher values of *m* compared to the two other clusters.

### 3.2 Cluster delineation reflects different demographic scenarios

Cluster delineation, and thus the magnitude of genetic changes, reflects and distinguishes three demographic scenarios.

As seen above, the most influential parameter for cluster delineation is *propR*. As this parameter determines the maximal population sizes on R and S, it results in differences in population dynamics among clusters (Figure 5). The mean compartment size of S is higher for Cluster 1, irrespective of the life cycle, which leads to higher initial population sizes and less genetic changes through time on S.

**Figure 5:**
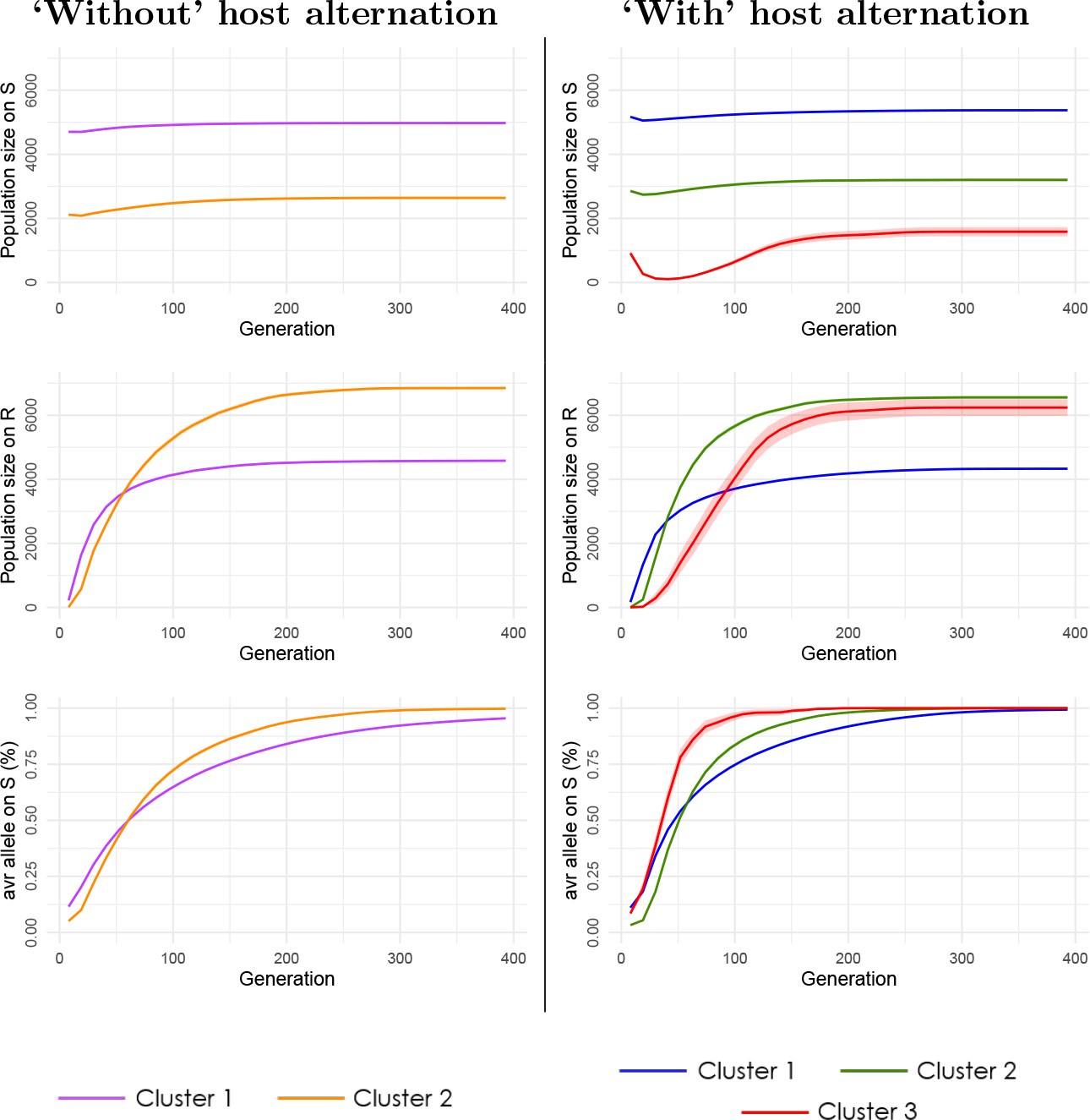
Temporal evolution of population sizes and virulent allele frequency depending on the cluster. The plotted results correspond to the mean temporal dynamics for all simulations in the corresponding cluster, among all simulations of the random simulation design. Shaded colour bands correspond to standard error intervals.

For both life cycles, the second most influential parameter is the initial frequency of avirulent allele (*f_avr_*), which plays a major role in the assignments to Cluster 2, with an over-representation of simulations with low values of *f_avr_* (Figures 4, S3). These low values of *f_avr_* result in fewer virulent individuals. It causes founder effects on R (Figure 5) and leads to more pronounced genetic changes through time (Figure 3).

A peculiar range of parameter values defines Cluster 3 observed ‘with’ host alternation. It results in evolutionary rescue dynamics. The low growth rate associated with a restricted size of S (high *propR*) causes an initial decrease in population size until near extinction (Figure 5). Then, the emergence of virulent individuals leads to the establishment of a new population on R and these individuals, by migrating back to S, prevent population extinction (see example in Figure S4, Cluster 3). Proportions of evolutionary rescue events differ significantly among clusters, with Cluster 3 composed almost entirely of such simulations (Table S3).

### 3.3 Inter- and intra-population genetic changes

To compare inter- and intra-population genetic changes associated with resistance overcoming, we focus on 1) mean dynamics (*i.e.* mean temporal evolutionary trajectories of population genetic indices on each cluster), and 2) realised dynamics (*i.e.* temporal evolutionary trajectories of population genetic indices for medoid or null model simulations of each cluster). Both mean and realised dynamics were computed from the variation at neutral loci and displayed using seven population genetic indices (Table 2).

#### Inter-population genetic signatures

All clusters show a peak in the mean dynamics of genetic differentiation between populations on R and S (*F_ST_ R − S*, Figure 2). This peak in differentiation occurs within the first 50 generations and is concomitant with resistance overcoming (Figure S5). The peak is more narrow in time ‘with’ host alternation, especially for Cluster 1, and the genetic differentiation is observable for a shorter period ‘with’ host alternation than in cases ‘without’ host alternation. The peak is higher for Cluster 2 than Cluster 1, which reflects stronger founder effects resulting from the settlement on R. ‘With’ host alternation, this peak in differentiation is magnified in Cluster 3 by the conjunction of the founder effect and the strong genetic drift on S resulting from the initial decrease in population size on S before the settlement on R leading to evolutionary rescue (Figure 5).

Without selection and for all simulations, the temporal *F_ST_* on S increases linearly through time at a slope depending on the relative importance of mutation and genetic drift forces (realised dynamics of temporal *F_ST_* for the null model, Figure S6). As the mutation rate is fixed, this slope depends on population size only. With selection, for all clusters we observe a two-phase dynamics of temporal differentiation on S (Figures 2, S6). The first phase corresponds to an initial increase in temporal *F_ST_*, stronger than without selection, that coincides with resistance overcoming. Temporal *F_ST_* reaches a maximum between 50 and 200 generations, depending on the cluster and the life cycle. This increase in temporal *F_ST_* on S is delayed but similar to the temporal *F_ST_* on R (see the example of the temporal *F_ST_* on R for Cluster 2 ‘with’ host alternation, Figure S7). The second phase exhibits more stable values of temporal *F_ST_*(slight increase or decrease). This second phase is concomitant with the regain in genetic diversity on both compartments, which favours homogenisation of allele frequencies distorted by the founder event (Figure 2). ‘With’ host alternation, simulations displaying strong genetic differentiation (realised dynamics of Cluster 2 and Cluster 3) show a steeper peak of differentiation (Figure S6) compared to ‘without’ host alternation. This is in accordance with the rapid decrease in differentiation between populations on R and S ‘with’ host alternation, which shifts the temporal *F_ST_* to the next phase of the dynamics (Figure S5).

#### Intra-population genetic signatures

For all clusters, we observe a two-phase dynamic of temporal change in gene diversity (expected heterozygosity, *H_E_*) on S. The first phase corresponds to a strong decrease in *H_E_*, until a minimum value is reached between 50 and 200 generations, followed by a slower increase towards a new mutation-drift equilibrium (Figure 2). However, the timing of this change differs among clusters. For Cluster 1 and Cluster 2 and both life cycles, the decrease in *H_E_* follows the generation of resistance overcoming, while the null model dynamics show no variations in *H_E_* (Figure S6). The decrease in gene diversity results from the immigration of less diverse pathogen individuals from the founding population on R that overcame the resistance. For Cluster 3 (‘with’ host alternation), a strong decrease in *H_E_* is observed for both the null model and the medoid realised dynamics, and the drop in *H_E_* precedes resistance overcoming. Unlike other clusters, this drop in *H_E_* is preceded for Cluster 3 by a strong decrease in the mean number of alleles *Mean LA* (Figures 2, S6), indicating a bottleneck.

Simulations display a peak in linkage disequilibrium, with a maximum value of *r̄_D_* being reached in the first 100 generations (Figures 2, S6). The variation in *r̄_D_* is to be examined in relation to the variation in *F_IS_*, which patterns differ between clusters and life cycles. ‘Without’ host alternation, *r̄_D_* and *F_IS_* display similar variations but slightly delayed in time, with a maximum value of *F_IS_* reached in the first 50 generations. This indicates admixture of genetically differentiated individuals on S. ‘With’ host alternation, *F_IS_* is null or displays slightly negative values for Cluster 1 and Cluster 2 while *r̄_D_* remains positive. This indicates that the signature of the admixture on S is rapidly being erased ‘with’ host alternation. In Cluster 3 ‘with’ host alternation, the peak in *r̄_D_* coincides with very negative values of *F_IS_*. The negative values in *F_IS_* results from the decrease in *H_E_* that is preceding the decrease in *H_O_* (Figures S6, S8 and see discussion 4.3).

## 4 Discussion

### 4.1 Genetic signatures of resistance overcoming

Disease outbreaks caused by pathogens impact both natural and human-managed ecosystems (*e.g.* agrosystems) (Anderson et al., 2004; Tobin, 2015; Savary et al., 2019). The number of emerging diseases is increasing exponentially and unprecedentedly during the last decades (Fisher et al., 2012). Understanding pathogen evolution is essential to comprehend how they affect ecosystems (Fischhoff et al., 2020) and to establish relevant disease management programs (Bonneaud and Longdon, 2020). Yet, this task is particularly challenging due to the rapid adaptation of pathogen populations (McDonald and Linde, 2002; Saubin et al., 2023a), and the high stochasticity of pathogen evolutionary trajectories (Parsons et al., 2018). A clustering method dedicated to time-series variations allows us to draw a typology of scenarios of eco-evolutionary dynamics associated with a strong selective event. We apply this model to a resistance overcoming event underpinned by static host compartments. This model and our findings can be extended to any system where pathogen populations evolve on different resources whose type and abundance do not change over time.

All the recorded population genetic indices are impacted by pathogen adaptation. Overall, resistance overcoming leads to a founder effect on the resistant host, with a differentiated sub-sampled population settling and growing on resistant hosts. Migrations between susceptible (S) and resistant (R) hosts then homogenise the genetic coancestry over all pathogen populations at a pace that depends on the migration rate and pathogen population sizes. Overcoming the plant resistance has a strong impact on the pathogen population genetic structure on susceptible hosts, with 1) a decrease in pathogen genetic diversity, 2) a peak in linkage disequilibrium, 3) a strong increase in temporal genetic differentiation between the initial and evolved populations, and 4) a peak in population differentiation between the susceptible and resistant compartments. The comparison of the evolution of the genetic indices through time, with and without selection, shows that these changes are signatures of evolution under selection and do not result from genetic drift only. Our first important result is thus that changes in population genetics of neutral markers allow identifying a selective event of resistance overcoming. We note that most of these genetic changes are transient, with a signature of resistance overcoming that vanishes in a few years only.

### 4.2 Typology of dynamics under resistance overcoming

#### Each evolutionary scenario represents distinct genetic signatures of the demographic outcomes of adaptation

Simulations can be distinguished by the magnitude of their genetic signatures and grouped into clusters that are indicative of different evolutionary scenarios. This clustering is strongly linked to variations in pathogen population sizes. Cluster 1 regroups simulations with the slightest genetic signatures, associated with a steady and slow demographic expansion. Hence, a large part of the simulations leads to signatures of particularly low magnitudes. Cluster 2 regroups simulations with stronger genetic signatures, associated with larger demographic expansions on the resistant compartment. In extreme cases ‘with’ host alternation, the few simulations assigned to Cluster 3 present the strongest genetic signatures, mainly associated with a specific demographic scenario, namely evolutionary rescue. These simulations are characterised not only by a strong demographic expansion on the resistant compartment (as for Cluster 2), but also by a demographic recovery on the susceptible compartment. Overall, founder effects lead to a demographic expansion that is all the more important that the resistant host is abundant, because of the logistic population growth. During the first generations following the settlement on the resistant compartment, the successive clonal reproduction events lead to a large population of few genotypes largely repeated, thus strongly entangling demographic variations and genetic changes.

#### Differences due to input parameters

In this model, the main determinant of pathogen population sizes is the proportion of resistant hosts in the landscape, and not the intrinsic demographic parameters (pathogen growth and migration rates). For both life cycles, the proportion of resistant hosts determines the initial pathogen population size and the size of the compartment available through adaptation. As such, it drives the strength of both population expansion and selection pressure exerted on the pathogen population. ‘With’ host alternation, this proportion also shapes the population demography in determining the likelihood of evolutionary rescue (See Figure 4 in Saubin et al., 2021). Pathogen control often leads to high proportions of resistant hosts in agricultural landscapes (Stukenbrock and McDonald, 2008; Zhan et al., 2015). Here we show that higher proportions of resistant hosts lead to more pronounced genetic changes in the pathogen population that overcome resistance.

The initial proportion of virulent alleles in the pathogen population is also a strong driver of the strength of the genetic signatures. This extent of standing genetic variation determines the proportion of individuals that will be able to respond to selection, in other words, the adaptive potential of the initial population. In particular, the proportion of virulent alleles impacts the number of individuals that settle the pathogen population on the resistant host, hence the genetic diversity of the founded population. A small proportion leads to fewer virulent individuals, and therefore a less diverse and more differentiated population founded on resistant hosts, as observed for frequent turnover of extinction and recolonisation (McCauley, 1991).

#### Differences due to the pathogen life cycle

‘Without’ host alternation, genetic signatures remain detectable for a longer period of time, because of sustained genetic admixture. Under this life cycle, gene flow results from the movements of a limited number of individuals determined by the migration rate. Migration occurs at each generation, which leads to a regular but progressive homogenisation of pathogen populations evolving on resistant and susceptible hosts. This accounts for the delay between the maximum differentiation observed between compartments at the time of the founder event, and the return to a low differentiation after the homogenisation of the populations. At the time of maximum differentiation, the immigration on susceptible hosts of individuals from the newly founded population on resistant hosts causes a Wahlund effect (Wahlund, 1928), that is a distortion of allele frequencies caused by the admixture of genotypes originating from different subpopulations. This explains the positive values of both the inbreeding coefficient and the linkage disequilibrium.

‘With’ host alternation, genetic differentiation between compartments fades more rapidly because of the obligate mating event taking place in a common alternate host, where alleles at each locus are reshuffled through sexual reproduction. It erases the Wahlund effect observed ‘without’ host alternation as a single event of sexual reproduction among all individuals is sufficient for a return to Hardy-Weinberg proportions (Rouger et al., 2016). This explains the small or slightly negative values of the inbreeding coefficient. Yet, linkage disequilibrium remains positive because the associations of alleles across loci are still preserved for some time, as recombination occurs within individuals (whose allele frequencies are inherited from either population). Last, all individuals are redistributed randomly between the two compartments. The death of avirulent individuals on resistant hosts distorts allele frequencies and regenerates differentiation between pathogens evolving on resistant and susceptible hosts. This life cycle increases gene flow and leads to a fast homogenisation of pathogen populations.

These differences between life cycles explain the observed differences in demographic variations and genetic changes in pathogen populations. Examples of these different genetic outcomes can be found in the literature: from conservation of genetic structure ‘without’ host alternation (*e.g.* in Leroy et al., 2013) to strong selective sweep and gene swamping ‘with’ host alternation (*e.g.* in Persoons et al., 2017). In addition to the plant pathogens on which our examples are based, this model could be applied more widely to other organisms with similar life cycles (*e.g.* agricultural pests such as aphids, Moran, 1992). However, to our knowledge, there is a lack of empirical studies (and adequate datasets) on these species.

#### Stochasticity and genetic signatures

Beyond the influence of input parameters and life cycle on demographic variations and genetic changes, a complementary analysis based on simulation replicates highlights that identical combinations of input parameters can lead to different outcomes (Appendix B). The generation at which the first virulent individuals actually settle on the resistant host drives the resulting demography and genetic structure. Moreover, the stochasticity impacts not only the timing of settlement but also the number of successful migration events between compartments, hence the genetic diversity of the founded population. As the virulent allele is recessive, it is more vulnerable to extinction ‘with’ host alternation (Saubin et al., 2021). Here we show that in addition to the stochasticity in the fate of the virulent allele, the life cycle ‘with’ host alternation also increases the stochasticity in the genetic signatures of resistance overcoming.

### 4.3 Genetic signatures characteristic of evolutionary rescue

Different processes can lead to the survival of the pathogen population even if its decline is approaching extinction. Three forms of such population ‘rescue’ are commonly described (Carlson et al., 2014): 1) the demographic rescue, when the population survival is only attributed to the increase in population size due to immigration of new individuals (Brown and Kodric-Brown, 1977), 2) the genetic rescue, when the survival of the population is attributed to the novel genetic variation brought by the immigration of new individuals, in a small population suffering genetic load (Thrall, 1998), and 3) the evolutionary rescue, when adaptive evolution lead to population recovery from negative growth initiated by environmental change (Gonzalez et al., 2013; Bell, 2017). The two latter forms of rescue closely link demography and selection, whereby selection at one locus determines the demography of the populations, and thus the neutral variation across the genome. The probability of fixation of alleles is strongly impacted by changes in population size (Otto and Whitlock, 1997), with the effect of genetic drift accentuated by a reduction in population size. It is therefore all the more important to focus on the interplay between selection and genetic drift in a population with fluctuating size (Gokhale et al., 2013; Z^̌^ivkovíc et al., 2019) to weigh up the balance between deterministic and stochastic processes that drive the evolutionary trajectories of pathogen populations.

In this study, the observed population rescue can be considered as demographic or evolutionary, depending on the definition of the population. If we consider distinctly pathogen populations on susceptible and resistant hosts, the adaptation of virulent individuals leads to their settlement on the resistant host, hence to the survival of the population on the susceptible host. The survival of the population on the susceptible host corresponds to demographic rescue resulting from the immigration of adapted individuals from the resistant host. If we consider a single population encompassing all individuals evolving on both hosts, the survival of the population corresponds to an evolutionary rescue event, *via* the adaptation to the new environment (*i.e.* the newly deployed resistant hosts). Such evolutionary rescue events occur mostly for the life cycle ‘with’ host alternation (Saubin et al., 2021). This is because of additional mortality that originates once a year from the massive redistribution of individuals after the sexual reproduction, with the death of all avirulent individuals that migrate to resistant hosts. Besides the additional mortality, the massive redistribution also increases the probability that a virulent individual migrates to the resistant host, and hence increases the probability of evolutionary rescue (Saubin et al., 2021). Here we demonstrate that such events can lead to strong and typical genetic signatures at neutral loci, that define a specific cluster ‘with’ host alternation (Cluster 3) and can thus be uncovered using population genetic indices. These dynamics are characterised by a bottleneck with few possible genotypes combining the remaining alleles. This is evidenced by the changes in several indices, such as the drop in the mean number of alleles, followed by the drastic reduction in gene diversity and the increase in linkage disequilibrium (Cornuet and Luikart, 1996). Unexpectedly, this bottleneck comes along with a strikingly negative value of inbreeding coefficient. To understand this result, we investigate the cause of the discrepancy between expected and observed heterozygosity. This is due to the clonal reproduction events that maintain identical genotypes during the epidemic phase, hence the value of observed heterozygosity remains constant over generations, whereas expected heterozygosity steadily decreases because the population size is collapsing very fast. Note that as the sampling takes place at the end of the clonal phase, the difference between expected and observed heterozygosity is magnified by the small genotypic drift that happened between clonal lineages over the handful of clonal generations. Overall, even for such drastic demographic events, the resulting genetic signatures remain transients, and thus can only be captured by using time sample data in the appropriate time window.

### 4.4 Temporal changes of demogenetic signatures

Among indices, we observe different temporal dynamics of genetic signatures of resistance overcoming. The fastest changes are observed for the genetic differentiation between S and R and the mean inbreeding coefficient, during the 50 first generations (*i.e.* the first five years) after the resistance overcoming. Changes in linkage disequilibrium and mean number of alleles are slower and detected within the 100 first generations. Finally, the slowest changes (and less sharp temporal signatures) are observed for the expected heterozygosity and the temporal genetic differentiation, with most of the detectable signal occurring between generations 50 and 200. Overall, under the modelled population sizes, no peak of genetic signature occurs after generation 200, and only a residual signal remains on population genetic indices.

For indices reflecting disequilibrium induced by the founder effect (*F_ST_ R − S*, *r̄_D_*, *Mean F_IS_*), genetic changes stabilise rapidly, especially when gene flow is enhanced by a life cycle ‘with’ host alternation. For other indices (temporal *F_ST_*, *H_E_*, *Mean LA*), when it exists, the difference with the null model (*i.e.* without selection) persists for a much longer period (for at least 300 generations for *Mean LA* and until the end of the simulated period for temporal *F_ST_* and *H_E_*). Following rapid adaption, we thus observe a return to genetic equilibrium in two steps. The first step involves the homogenisation of allele frequencies both within and between populations thanks to the convergence to Hardy-Weinberg and migration-drift equilibria. This occurs very fast because the modelled system is regularly subjected to the stabilising effect of sexual reproduction (Rouger et al., 2016). We hypothesise that this return would be slower for systems that deviate from this mode of reproduction, either because of an increased rate of clonality (Reichel et al., 2016) or because of selfing (Jullien et al., 2019). The second step is the return to allele numbers and frequencies expected under the migration-mutation-drift equilibrium. This evolution is slower because it relies on the progressive change in allelic states and potential recovery of the lost genetic diversity that happened during the upheaval caused by resistance overcoming.

The timing of the modelled events may appear rapid but is consistent with empirical observations. For example, in the poplar rust pathogen (*Melampsora larici-populina*) which alternates on larch every year, the overcoming of resistance RMlp7 in poplars led in 1994 to a strong genetic disequilibrium. This was followed by a quick return to Hardy-Weinberg equilibrium the following year and drastic changes in the population genetic structure occurring in less than four years (Persoons et al., 2017; Louet et al., 2023).

Overall, studies employing time-series remain rare compared to the amount of work focusing on the genetic analysis of one contemporary pathogen population (Buffalo and Coop, 2019, 2020; Pavinato et al., 2021). Analyses of time-series are mostly used to analyse the speed and timing of selection for life history traits (Rouzic et al., 2011, 2015), loci under positive or fluctuating selection (Bergland et al., 2014; Foll et al., 2015), or coevolution between host and parasite (Decaestecker et al., 2007; Gandon et al., 2008; Blanquart and Gandon, 2013). We note that studies including neutral markers mainly use them, to date, to draw a statistical inference of loci under selection rather than to document the change in demography. In the rare cases where time-series are used to study specifically neutral genetic evolution, the data typically exhibit a limited temporal range (*e.g.* two time points, Pavinato et al., 2021). However, as temporal data allow to trace the changes in allele frequency through time, the analysis of neutral markers can improve our inference and understanding of evolutionary (Dehasque et al., 2020; Feder et al., 2021; Saubin et al., 2023a,b) and coevolutionary (Z^̌^ivkovíc et al., 2019) processes. Temporal full genome datasets available in *Drosophila melanogaster* (Bergland et al., 2014) also prompted new theoretical developments regarding the effect of seasonal population size changes and fluctuating selection on neutral variants (Wittmann et al., 2017, 2023). In cases of rapid adaptation, we specifically show here that the resulting genetic signatures may be very brief and require time samples around the selection event. This is in accordance with the study of Saubin et al. (2023b), in which several time samplings with rarefaction are tested and show high identifiability of the transient genetic signatures even if a strong thinning is applied to time-series data. Note that in the case of coevolution, and unlike our study, the host compartment is no longer static and should be monitored too (Brown and Tellier, 2011; Z^̌^ivkovíc et al., 2019) at the adequate temporal scale. Studies focusing on one or two time points may allow documenting only part of the coevolutionary dynamics and likely fail to highlight transient dynamics, which provide the most relevant information regarding the demogenetic interplay. Conversely, epidemiological considerations lead to focus on the time when the settlement is detected, therefore when the genetic signature is the strongest. These considerations tend to neglect the initial state, as well as the return to a new equilibrium, which may nevertheless occur in a relatively short time scale. A key result of our study is to demonstrate that neutral markers can be used to uncover demogenetic processes due to selection events (see also Z^̌^ivkovíc et al. (2019) for coevolution). As a validation, this approach has been successfully applied to temporal data, allowing to infer demographic scenarios and parameter values of a major event of resistance overcoming by the poplar rust pathogen (Saubin et al., 2023b).

## Acknowledgements

We thank Alexis Sarda for insightful answers on the Dtwclust package, Raphäel Leblois for the perceptive discussion that greatly helped us to formalise the results, and Lydia Bousset, Virginie Ravigńe, and Maria Orive for constructive comments of a previous version of this manuscript.

## Data accessibility

R scripts for clustering of time-series and statistical analyses will be made available on a public GitLab repository at the time of publication. An executable file will be provided on this repository to run the population genetics simulations.

## Author contributions

MS, SS, AT, and FH conceived and designed the study. MS and SS produced the code and ran the simulations. MS and FH analysed the data and prepared the manuscript. All authors revised and approved the manuscript.

