## Appendix for "Neutral genetic structuring of pathogen populations during rapid adaptation"

**Corresponding author:**

Fabien Halkett

### A Model description

We provide here a detailed description of the reproduction and migration events of the model, as presented in previous works based on a similar model (Saubin et al. 2021, 2023b).

#### A.1 Reproduction events

Each discrete generation corresponds to a reproduction event, either sexual or clonal. At each reproduction event, successive generations do not overlap so that parents give way to offsprings and the new population is composed exclusively of the newly produced individuals. The within-compartment dynamics of the pathogen population are provided by the following equations:

For the sexual reproduction event, the population size is considered constant before and after the reproduction event:

$$N_{n+1} = N_n \tag{1}$$

With $N_{n+1}$ the population size at generation $n+1$ and $N_n$ the population size at generation $n$. Sexual offspring genotypes are obtained by random association of gametes produced by the parental population.

For each clonal reproduction event on S, R, or A, the population growth has a logistic function, with the regulation that applies locally:

$$N_{n+1} = N_n + (r-1) \times N_n \times (1 - \frac{N_n}{K_i}) \tag{2}$$

With $r$ the growth rate of the pathogen population, $K_i$ the carrying capacity of the compartment ($K_S$, $K_R$, or $K_A$ for S, R, or A respectively). For all clonal reproduction events, offspring genotypes are drawn randomly with replacement from the parental population in which all parental pathogen individuals have the same uniform probability to reproduce.

#### A.2 Migration events

A regular two-way migration event takes place each generation before the reproduction event, between R and S. The number of migrants is determined by a fixed migration rate ($m$) multiplied by the number of individuals on the source compartment. In this effective migration scheme, migrants are always successfully established on the destination compartment, even if the number of individuals on this compartment reached its maximum carrying capacity. Thereby, this choice enables the immigration of new pathogens regardless of the size of the population, as it is observed in natural populations for plant pathogens for example.

For the life cycle ‘with’ alternation, the annual sexual reproduction event coincides with the obligate migration of the entire pathogen population to and from the alternate compartment. The first migration event takes place once per year after $g-2$ clonal reproduction events on R and S. For this migration event, an established proportion of individuals $\tau$ is picked randomly from R and S and migrates to A. R and S compartments are then emptied of any pathogen for the next two generations. After the two reproduction events (sexual and clonal) on A, the second migration event redistributes randomly all individuals from A to R and S, in proportion to the compartment carrying capacities ($K_S$ and $K_R$), leaving no living individuals on A.

### B Regular simulation design and stochasticity in the genetic signatures

To complement results from the random simulation design and to evaluate the effect of stochasticity on the variability of evolutionary trajectories, we perform a regular simulation design composed of 100 replicates of 216 fixed combinations of parameter values (Table S1).

Table S1: Input parameters and their range of variations for the regular simulation design. One hundred replicates are run for each combination of parameters, and each simulation is run for 400 generations.

| Variable | Description | Levels |
|---|---|---|
| $f_{avr}$ | Initial frequency of the *avr* allele in the pathogen's population | **4 levels**: 0.01; 0.03; 0.1; 0.32 |
| $m$ | Migration rate between R and S compartments | **3 levels**: 0.001; 0.01; 0.1 |
| $r$ | Growth rate of the pathogen | **3 levels** : 1.2; 1.6; 2.0 |
| $propR$ | Proportion of resistant hosts in the landscape | **3 levels**: 0.1; 0.5; 0.9 |
| $Cycle$ | Life cycle of the pathogen | **2 levels**: 'without' host alternation; 'with' host alternation |

As for the random simulation design, we perform clustering analyses 'with' and 'without' host alternation to identify the proportion of simulations that result from the same combination of input parameters (replicates) but are assigned to different clusters (Table S2).

Table S2: Cluster description for the clustering based on the regular simulation design. This table contains the number of simulations assigned to each cluster, and the percentages of simulations from the same set of parameters (100 replicates) which have been assigned to different clusters ('Different assignments (%)'). Only simulations leading to resistance overcoming have been kept for the clustering analysis.

| Cycle | Cluster | Number of simulations | Different assignments (%) |
|---|---|---|---|
| 'Without' host alternation | 1 | 2966 | 5.9 |
| | 2 | 2605 | 12.4 |
| 'With' host alternation | 1 | 6937 | 16.9 |
| | 2 | 1332 | 85.0 |
| | 3 | 42 | 97.6 |

As for the clustering based on the random simulation, temporal changes in population genetic indices from the regular simulation design group into two clusters 'without' host alternation, and three clusters 'with' host alternation (Table S2). For both life cycles, Cluster 1 regroups the majority of the simulations (53 % and 83 % of the simulations 'without' and 'with' host alternation, respectively, Table S2). Notably, we identify simulation replicates (*i.e.* with identical combinations of parameter values) that show different outcomes in the temporal changes of population genetic indices. Hence, simulation replicates can be assigned to different clusters, demonstrating a substantial degree of stochasticity. These replicates assigned to different clusters represent 6% and 12% of the simulations in Cluster 1 and Cluster 2 'without' host alternation, respectively, and 17%, 85% and 98% of the simulations in Cluster 1, Cluster 2, and Cluster 3 'with' host alternation, respectively (Table S2). Therefore, cluster delineation is not defined by specific combinations of parameter values, but rather by the inherent demographic scenarios underpinning the genetic dynamics.

This variation among simulations replicates results from the inherent stochasticity of the model. In particular, migrants are picked randomly. The generation at which the first virulent individuals actually settle on the resistant host drives the resulting demography and genetic structure. One year is composed of one sexual generation and ten clonal generations. Therefore, if the settlement occurs right after the sexual generation, several clonal generations follow, building a strongly differentiated population on resistant hosts. Conversely, if the settlement occurs at the end of the season of clonal reproductions, the population size does not change much before the next sexual reproduction reshuffles alleles in the population. Moreover, the stochasticity impacts not only the timing of settlement but also the number of successful migration events between compartments, hence the genetic diversity of the founded population. Beyond the inherent properties of the model, the life cycle 'with' host alternation also adds stochasticity. 'Without' host alternation, the first arrival of a pathogen individual on the resistant compartment settles a population, hence leading inevitably to resistance overcoming. Conversely, 'with' host alternation, a first arrival is necessary but not sufficient for a sustained settlement. The regular emptying of compartments before sexual reproduction and random individual redistribution can lead thereafter to stochastic settlements on resistant hosts as long as the virulent allele is rare (*e.g.* occupied compartment one year and cleared compartment the next year). As the virulent allele is recessive, it is more vulnerable to extinction 'with' host alternation (Saubin et al., 2021).

### C Supplementary tables and figures

Table S3: Cluster description for the clustering based on the random simulation design. This table contains the number of simulations assigned to each cluster, and the percentages of simulations leading to evolutionary rescue. Only simulations leading to resistance overcoming have been kept for the clustering analysis. The $P-value$ corresponds to the $P-value$ of the exact test of Fisher to determine if differences in proportions of evolutionary rescue events among clusters are significant.

| Cycle | Cluster | Number of simulations | Evolutionary rescue (%) | Fisher $P-value$ |
|---|---|---|---|---|
| 'Without' host alternation | 1 | 4067 | 1.3 | $4.6 \times 10^{-5}$ |
| | 2 | 1575 | 3.0 | |
| 'With' host alternation | 1 | 5858 | 3.7 | $< 2.2 \times 10^{-16}$ |
| | 2 | 1581 | 10.6 | |
| | 3 | 81 | 97.5 | |

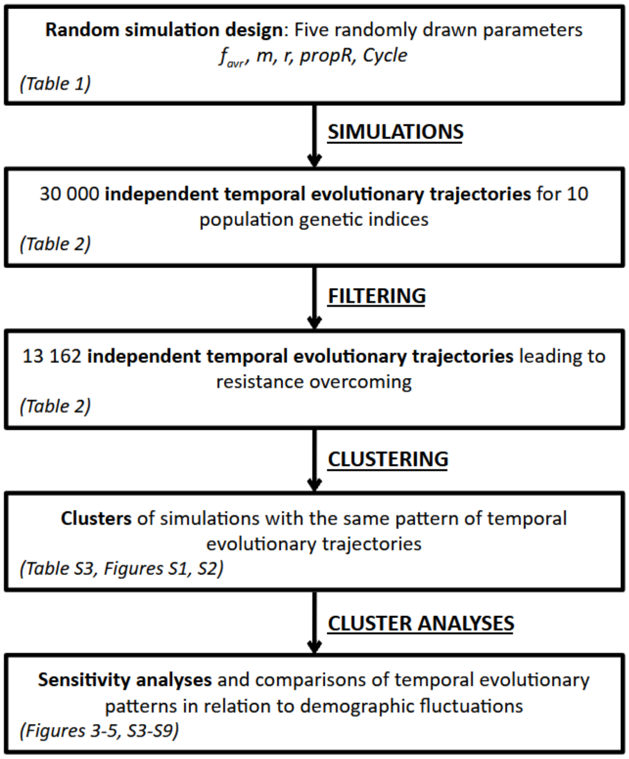


Figure S1: Main steps of the method, from the simulation design to the results analyses. Each step is associated with the related Tables and Figures.

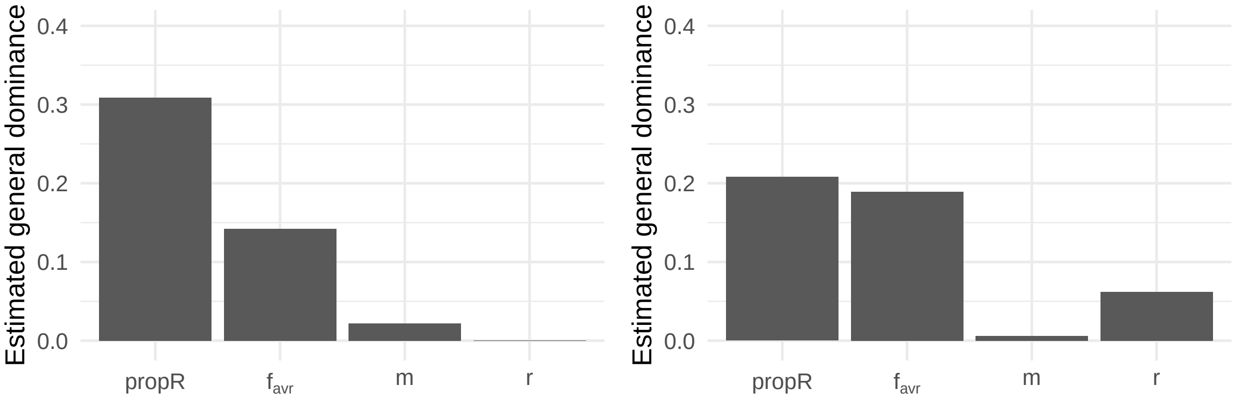


(a) 'Without' host alternation

(b) 'With' host alternation

Figure S2: Estimated general dominance calculated from a multinomial logistic regression applied to the cluster assignment for each simulation of the random design. The general dominance was estimated for the four input parameters: $propR$, $f_{avr}$, $m$, and $r$.

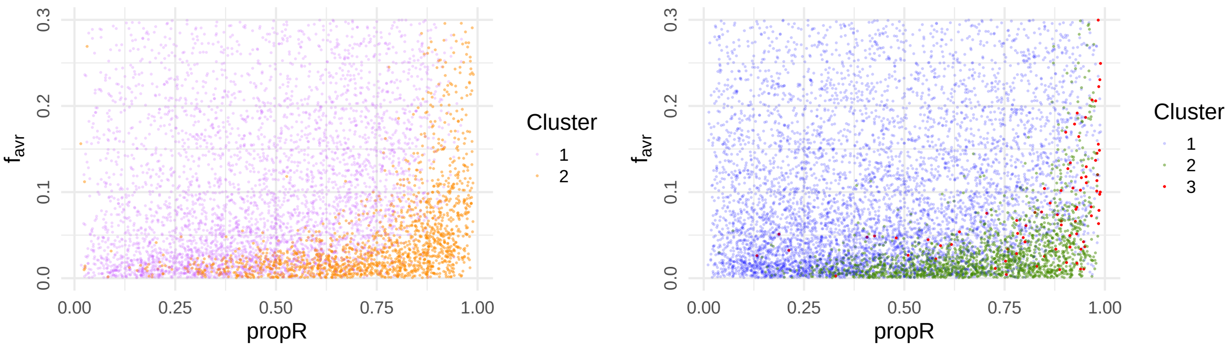


(a) 'Without' host alternation

(b) 'With' host alternation

Figure S3: Distribution of $propR$ and $f_{avr}$ input parameters for each simulations of both life cycles. Each point represents a simulation of the random simulation design that led to resistance overcoming, coloured by its cluster assignment.

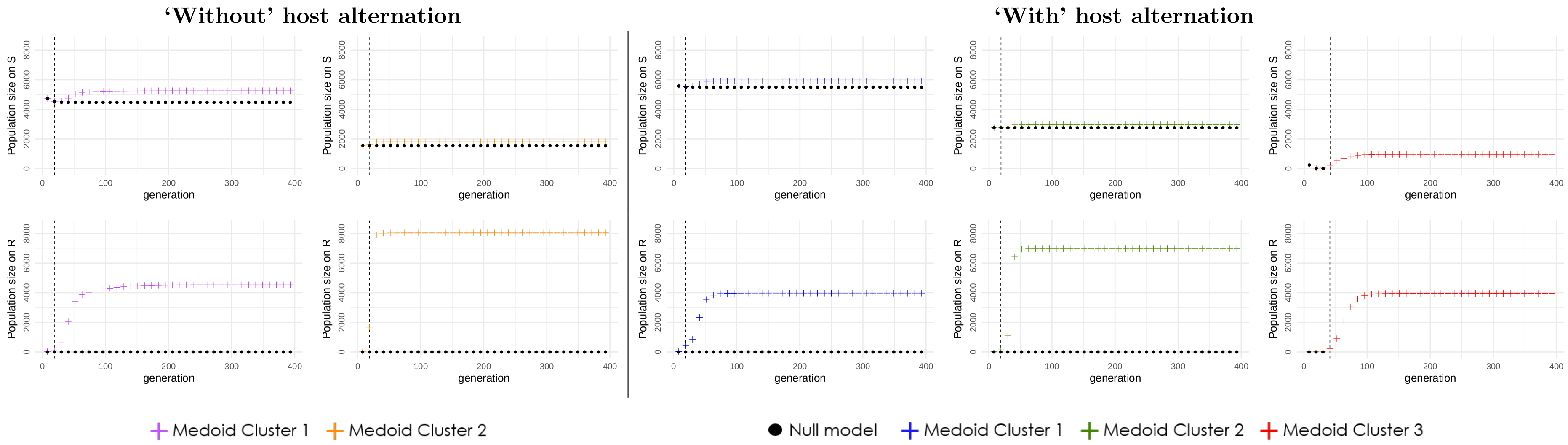


Figure S4: Comparison of the temporal evolution of population sizes, between medoid simulations and null models. The plotted results correspond to the medoid dynamic of each cluster, with ('Medoid', cross shapes) or without ('Null model', round shapes) virulent allele, that is, with or without resistance overcoming.

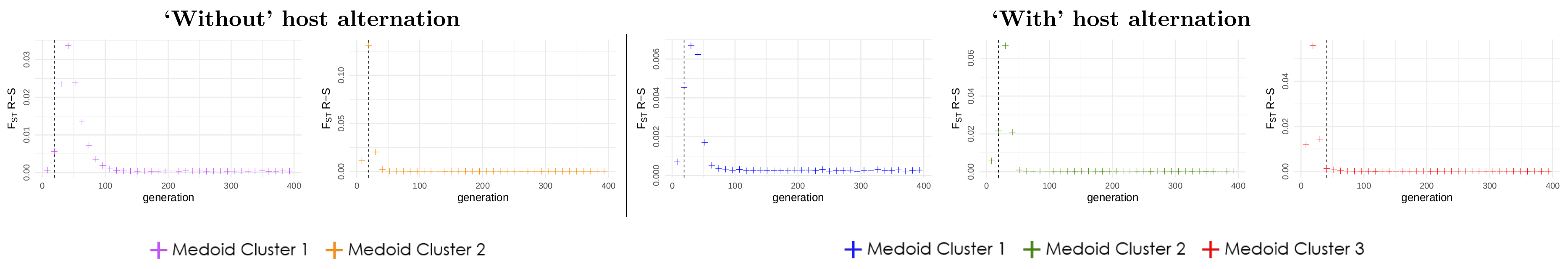


Figure S5: Temporal evolution of $F_{ST}$ $R - S$. The plotted results correspond to the medoid dynamics for each cluster and each life cycle. For each of the plotted medoids, the vertical dashed line corresponds to the generation of resistance overcoming.

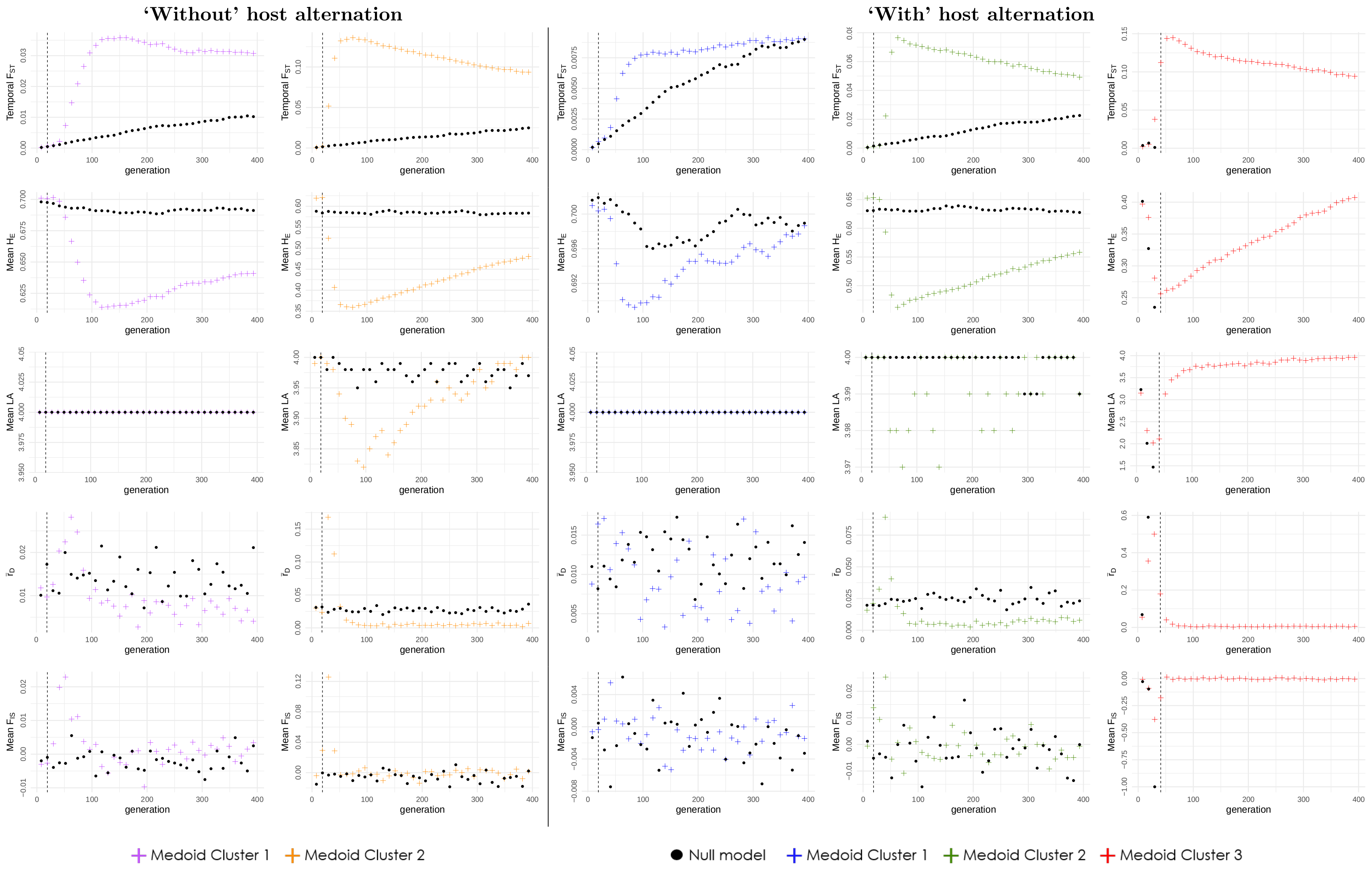


Figure S6: Comparison of the temporal evolution of population genetic indices, between medoid simulations and null models. The plotted results correspond to the medoid dynamic of each cluster, with ('Medoid', cross shapes) or without ('Null model', round shapes) introducing virulent allele, that is, with or without resistance overcoming. For each of the plotted medoids, the vertical dashed line corresponds to the generation of resistance overcoming. Cluster 3 'with' host alternation displays no round shapes after the generation of resistance overcoming because the population goes extinct in the 'Null model'.

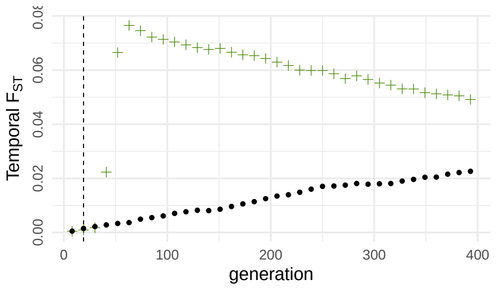


(a)

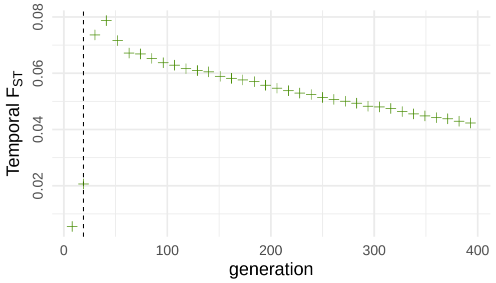


(b)

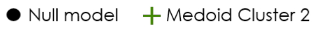


Figure S7: Comparison of the medoid dynamics of *Temporal* $F_{ST}$ on (a) S and (b) R for Cluster 2 'with' host alternation. Subfigure (a) is already presented in the main document (Figure S6, 'with' host alternation, Cluster 2).

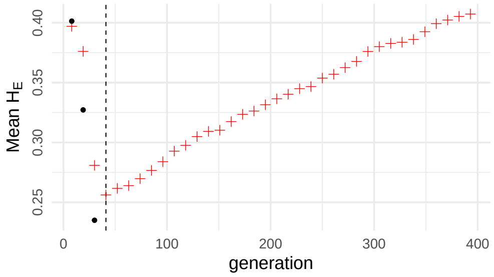


(a)

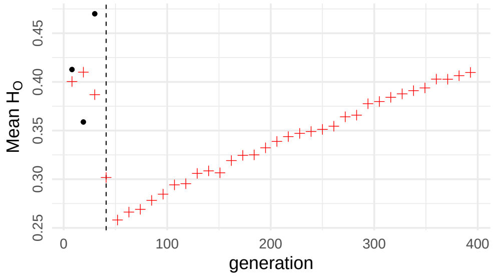


(b)

● Null model + Medoid Cluster 3

Figure S8: Comparison of the medoid dynamics of (a) $Mean\, H_E$ and (b) $Mean\, H_O$ on S for Cluster 3 'with' host alternation. Subfigure (a) is already presented in the main document (Figure S6, 'with' host alternation, Cluster 3).
